# Activity-dependent constraints on catecholamine signaling

**DOI:** 10.1101/2023.03.30.534970

**Authors:** Li Li, Akshay Rana, Esther M. Li, Jiesi Feng, Yulong Li, Michael R. Bruchas

**Author notes:** Co-corresponding authors. Li, Li, MD, PhD and Michael R. Bruchas, Ph.D, and respectively. Equal contribution (Akshay Rana, Esther M. Li).

## Abstract

Catecholamine signaling is thought to modulate cognition in an inverted-U relationship, but the mechanisms are unclear. We measured norepinephrine and dopamine release, postsynaptic calcium responses, and interactions between tonic and phasic firing modes under various stimuli and conditions. High tonic activity *in vivo* depleted catecholamine stores, desensitized postsynaptic responses, and decreased phasic transmission. Together this provides a clearer understanding of the inverted-U relationship, offering insights into psychiatric disorders and neurodegenerative diseases with impaired catecholamine signaling.

## Introduction

Norepinephrine (NE) and dopamine (DA) act as major catecholaminergic neuromodulators which act on G-protein coupled receptors (GPCRs) to regulate arousal states^1–3^, attention^4–6^, learning & memory ^5, 7^, and sensory-motor functions^8–10^. Interestingly, the activity of the locus coeruleus (LC), the brainstem nucleus that serves as the principal source of cortical NE, has been proposed to exhibit an inverted-U relation with task performance^8^. Similarly, dopamine D1 receptor activation exhibits an inverted-U relation with working memory^7^; and the activity of DA neurons in the ventral tegmental area (VTA) produces an inverted-U relation with short-term memory performance^11^. While the positive dose-response relation at lower neuronal firing is unsurprising, our mechanistic understanding of decreased cognitive function at the upper end of the catecholaminergic action remains incomplete. It has been hypothesized that these two neuromodulators may deplete at the upper limit of firing for the respective catecholaminergic neurons, but this hypothesis has not been tested *in vivo*, due to a lack of high-resolution tools to selectively detect NE and DA release. Moreover, it is unclear whether the phasic mode of transmission (in LC 10-20Hz, 200-500ms^12^, and in VTA ∼20Hz, <200ms^13, 14^) is affected by higher tonic firing. The recent introduction of genetically encoded fluorescent sensors for NE^15^ and DA^16, 17^ can help to address these questions by overcoming the sensitivity, speed, and spatial limitations of cyclic voltammetry^18^. Here, we report that high tonic LC^NE^ activity quickly depletes NE, decreases the postsynaptic excitability response, and reduces NE release to subsequent phasic firing events *in vivo*. Similarly, high tonic VTA activity can also deplete DA and reduce DA release to phasic firing *in vivo*. Moreover, acute stress affects primarily the postsynaptic response and likewise reduces phasic NE release. Thus, these findings implicate a possible synaptic mechanism in which catecholamine signaling becomes progressively impaired at higher neuronal firing due to depleted catecholamine stores and desensitized postsynaptic responses, facilitating a better understanding of the non-linear behavioral effects of catecholamine neuromodulation from the non-linear effects in catecholamine-induced GPCR signaling.

## Results

To examine how noradrenergic signaling modulates arousal states via downstream effector regions, we chose two projections of LC^NE^ neurons to well-known wake-promoting structures, one to the posterior basal forebrain (BF) and the other to the medial thalamus (Thal) ^19, 20^. As shown in **Fig 1a** we virally expressed a cre-inducible fluorescent yellow protein (AAV5-EF1α-DIO-EYFP) in the LC^NE^ neurons of Dbh-cre mice, and visualized robust labeling of LC axons in the BF and Thal. Next, we measured axonal NE release *in vivo* by expressing a genetically encoded fluorescent NE sensor (GRAB_NE2m) in BF neurons, and a cre-inducible optogenetic actuator (ChrimsonR) in the LC^NE^ neurons of Dbh-cre mice; and implanted an optical fiber in the BF for both fiber photometry and photostimulation experiments (**Fig 1b**).

**Figure 1.**
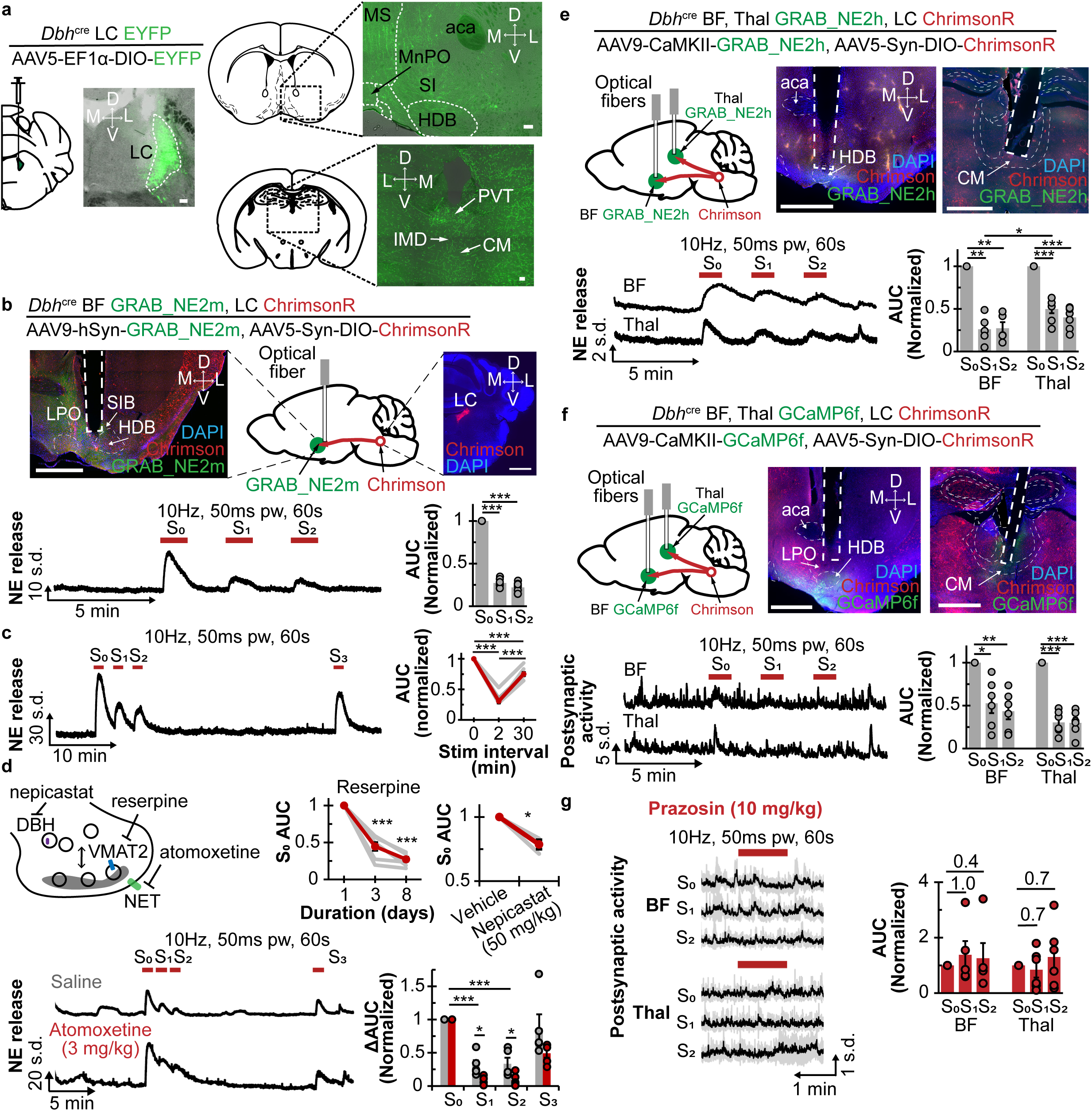
**a.** Left, schematic shows injection of cre-inducible EYFP virus in the LC of Dbh-cre mice, accompanied by coronal sections showing EYFP expression in the LC. Right, schematic and coronal sections showing the basal forebrain (top) and medial thalamus (bottom). Scale bars are 100µm. aca, anterior. CM, centromedian. HDB, horizontal limb of the diagonal band of Broca. IMD, intermediodorsal nucleus. MS, medial septum. MnPO, median preoptic nucleus. PVT, paraventricular thalamus. SI, substantia innominata. **b.** Top, schematic and coronal sections show expression of ChrimsonR in LC^NE^ neurons, GRAB_NE2m in the basal forebrain (BF), and an implanted optical fiber in the BF. Scale bars are 1mm. HDB, horizontal limb of the diagonal band. LPO, lateral preoptic area. SIB basal portion of substantia innominata. Bottom left, trace shows NE release in response to repeated axonal photostimulation at 10Hz, 50ms pw for 60s at 2 min intervals. Bottom right, quantification of the area under curve (AUC) normalized to the first stimulation (N=8). ***p<0.001. **c.** Left, trace shows NE release with repeated photostimulation at 10Hz, 50ms pw for 60s at 2min and 30min intervals. Right, graph shows corresponding AUC normalized to the first stimulation (N=8). ***p<0.001. **d.** Top left, diagram shows the drugs used to alter presynaptic NE metabolism. Top right, graphs show normalized AUC S_0_ after multiple days of reserpine treatment (see **SFig 1C** for treatment course, N=7) and 2h after intraperitoneal (*i.p*.) administration of 50mg/kg nepicastat (N=3). Bottom left, traces of NE release 20min after *i.p*. saline (gray) or atomoxetine (red) with repeated 10Hz photostimulations. Bottom right, graph shows changes to AUC normalized to the first stimulation (N=3). *p<0.05, ***p<0.001. **e.** Top left, schematic shows expression of Chrimson in the LC^NE^ neurons and GRAB_NE2h in the BF and Thal, and implanted optical fibers in the two areas. Top right, histological coronal brain slices show viral expressions and optical fiber placements in the Thal and BF. Scale bars are 1mm. CM, centromedian nucleus. aca, anterior commissure. HDB, horizontal limb of diagonal band. Bottom left, traces show NE release in response to 10Hz photostimulation in the BF and Thal. Bottom right, plot shows corresponding AUC normalized to the first stimulation in the BF (N=5) and Thal (N=7), respectively. *p<0.05, **p<0.01, ***p<0.001. **f.** Top left, schematic shows expression of Chrimson in the LC^NE^ neurons and GCaMP6f in the BF and Thal, and implanted optical fibers in the two areas. Top middle and right, histological coronal brain slices show viral expressions and optical fiber placement in the Thal and BF. Scale bars are 1mm. CM, centromedian nucleus. aca, anterior commissure. HDB, horizontal limb of diagonal band. Bottom left, traces show postsynaptic activity in response to 10Hz photostimulation in BF and Thal. Bottom right, plot shows AUC normalized to the first stimulation in the BF (N=6) and Thal (N=6), respectively. *p<0.05, **p<0.01, ***p<0.001. **g.** Left, average traces with standard deviation show the postsynaptic activity in the BF and Thal in response to the repeated 10Hz photostimulations 20min after *i.p.* 10mg/kg prazosin injection. Right, plot shows AUC normalized to the first stimulation in BF (N=6) and Thal (N=6), respectively.

LC^NE^ tonic frequency typically ranges from 1-6Hz^12, 21^, though it has been shown to go up to ∼8-10Hz with significant stress^22, 23^. Given tonic 10Hz somatic LC^NE^ activation for 10min has previously been shown to decrease NE level in the medial prefrontal cortex^2^, we reasoned that 10Hz may provide a suitable starting frequency to examine NE depletion. When LC^BF^ axons were repeatedly photostimulated at 10Hz, 50ms pulse width (p.w.) for 60s at 2min intervals, the initial stimulation (S_0_) elicited a robust increase in NE release and the measured signal change is set as 100%, but the following stimulations (S_1_ and S_2_) elicited only 27±2% and 22±2%, respectively, of the initial NE release amount (**Fig 1b**, **supplemental Table 1**), consistent with NE depletion from axonal release sites. Repeating stimulation 30min after such a depleting stimulation induced 75±3% of the initial NE release (**Fig 1c**), consistent with NE repletion. Moreover, we observed that NE release to S_2_ is similar to S_1_, suggesting the existence of two NE pools: a fast- and a slow-repleting.

Next, we determined whether the proportions of these fast- and slow-repleting NE pools change by pharmacologically altering NE metabolism in the axon terminals. Using reserpine to inhibit vesicular monoamine transporter or nepicastat to inhibit dopamine β-hydroxylase, we found that both treatments significantly reduced NE release (**Fig 1d**), but did not alter the extent of NE release depletion and repletion (**SFig 1b,c**), indicating that the reduced NE is likely distributed equally amongst the fast- and slow-repleting pools. Interestingly, blocking NE reuptake transporter (NET) with atomoxetine (ATOM) when compared to the saline control (SAL), produced a significantly slower decay (τ_ATOM_ 0.006±0.002s versus τ_SAL_ 0.022±0.004s), but not rise, of NE fluorescence after stimulation (**SFig 1d**). We also found a significantly reduced NE release to S_1_ (SAL: 31±7% versus ATOM: 9±3%) and S_2_ (SAL: 33±9% versus ATOM: 9±4%) under the same pharmacological treatment conditions. These results indicate that the fast-repleting NE pools rely on NE reuptake, and that the observed reduction in NE fluorescence to S_1_ and S_2_ is not likely due to the internalization of GRAB_NE sensors, something previously established *in vitro*^15^. Furthermore, when compared to SAL control, selective agonism of α2-adrenergic receptors (α2-AR) signaling with dexmedetomidine (DEXMED) and blockade of α2-AR with atipamezole (ATI) significantly and bidirectionally altered NE release to S_1_ (SAL: 48±7% DEXMED: 22±4%; SAL 26±3%, ATI: 39±2%) and S_2_ (SAL: 37±5%, DEXMED: 21±3%; SAL 22±3%, ATI: 34±2%, **SFig 1e**). This result suggests that α2-mediated signaling can modulate the sizes of the fast- and slow-repleting NE pools, but only partially. As an additional control, we showed that 1 mg/kg ATI is able to antagonize the sedating effect of 50mcg/kg DEXMED (**SFig 1f**). These experiments show that although NE depletion cannot be wholly prevented, the extent of depletion can be altered by modifying NE metabolism.

To determine if the depletion-repletion properties observed in the BF are similar in other regions where NE is released, we next examined NE release in the (thalamic) LC^Thal^ projections using the GRAB_NE2h sensor (a version that has higher NE affinity than, but with comparable on-off kinetics to GRAB_NE2m) and LC^NE^-expressed ChrimsonR (**Fig 1e**). Using repeated photostimulation in BF showed significantly decreased NE release (S_1_ 31±6% and S_2_ 30±6% of S_0_) as seen before, suggesting that GRAB_NE2h and 2m consistently detect this NE depletion. Interestingly, the NE release induced in the Thal (S_1_ 53±11% and S_2_ 43±9% of S_0_) was significantly less than in the BF. The difference in induced NE release points to brain region-specific proportion of fast- and slow-repleting NE pools, potentially from the differing levels of NE transporter and α2-adrenergic receptors (AR) expressed on a given neuron^24^, which were shown to modulate these pool fractions.

We next determined whether local NE depletion affects local postsynaptic activity *in vivo*. Similar to the above strategy, ChrimsonR was again expressed in the LC^NE^ neurons of Dbh-cre mice, but the calcium sensor GCaMP6f was expressed locally in BF and Thal neurons (**Fig 1f**). When we repeated photostimulation of LC^NE^ axons at 10Hz (50ms pw) for 60s either in the BF or Thal, the local postsynaptic rise in calcium was significantly decreased with S_1_ (BF: 53±11%, Thal: 31±6% of S_0_) and S_2_ (BF: 43±9%, Thal: 30±6% of S_0_) stimulations, mirroring the depleted NE release at the axons. The increased postsynaptic calcium in response to stimulations of the LC^NE^ axons was blocked with prazosin (10 mg/kg), a selective α1 adrenergic receptor antagonist, but not by propranolol (10 mg/kg), a β adrenergic receptor antagonist (**Fig 1g, SFig 1g,h**), consistent with the well-established α_1_/G_q_-mediated signaling cascade that leads to increased intracellular calcium^25^. It also demonstrates that the postsynaptic rise in calcium dynamics is indeed primarily mediated by the NE release and not by other co-released transmitters or neuromodulators in these two LC projecting brain regions.

As LC^NE^ tonic frequency ranges from 1-6Hz^12, 21^, we then examined whether NE depletes at the higher frequencies of firing. Using mice with ChrimsonR expressed in the LC^NE^ and GRAB_NE2m expressed in the BF, we measured NE release during 30min of no stimulation versus axonal photostimulations at 3, 5, and 10Hz (10ms pw). As shown in **Fig 2a**, the stimulations elicited an initial, frequency-dependent increase in NE release, followed by a slow decline, consistent with NE depletion. When GRAB_NE2m was expressed in the Thal, a similar pattern of NE release kinetics was observed in a frequency-dependent stimulation of thalamic axons (**Fig 2b**). We also investigated the changes to postsynaptic activity in BF and Thal during 30min stimulations in a similar manner by using GCaMP6f to measure calcium dynamics. Despite the expected higher variability in postsynaptic activity, we observed a transient increase in postsynaptic activity in a frequency-dependent manner in both BF and Thal neurons (**SFig 2a**, **Fig 2c,d**), again mirroring the pattern of axonal NE release in these regions. These experiments indicate that NE depletion occurs with prolonged, elevated tonic activity.

**Figure 2.**
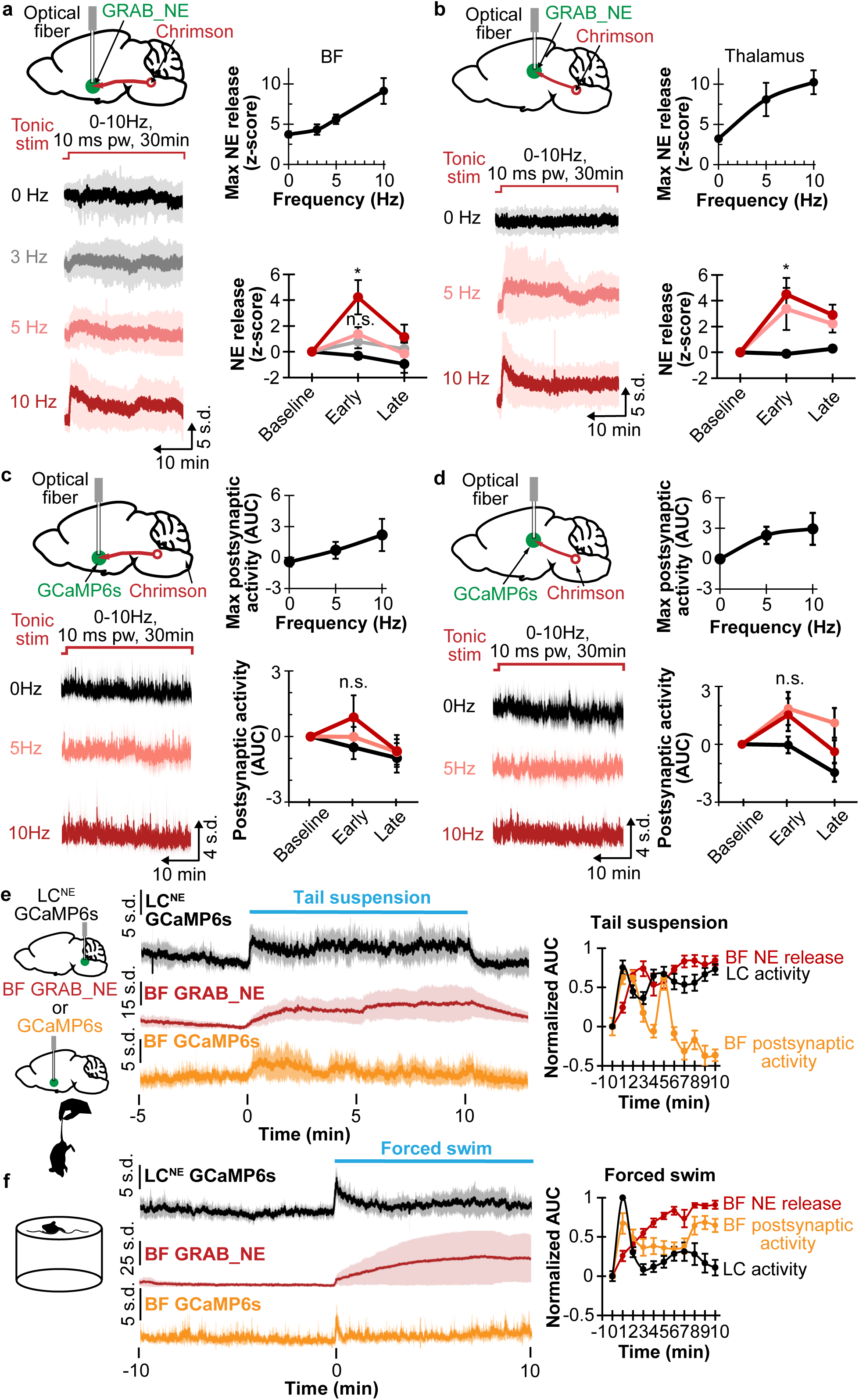
**a.** Left top, schematic shows Chrimson in the LC^NE^ neurons, GRAB_NE in the BF, and implanted optical fiber in BF. Left bottom, average traces with standard deviation show NE release with 30min photostimulation at 0, 3, 5, and 10Hz 10ms pw. Right top, plot shows maximum NE release in relation to stimulation frequency (N=8). Right bottom, plot shows NE release at baseline (1min before stimulation), early stimulation (1-5min), and late stimulation (25-30min) at different stimulation frequencies (N=8). *p<0.05. n.s. not significant. **b.** Left top, schematic shows Chrimson in LC^NE^ neurons, GRAB_NE in the Thal, and implanted optical fiber in Thal. Left bottom, average traces with standard deviation show NE release with 30min photostimulation at 0, 5, and 10Hz 10ms pw. Right top, plot shows maximum NE release in relation to stimulation frequency (N=7). Right bottom, plot shows NE release at the baseline (1min before stimulation), early stimulation (1-5min), and late stimulation (25-30min) at different stimulation frequencies (N=7). *p<0.05. **c.** Left top, schematic shows Chrimson in the LC^NE^ neurons, GCaMP6f in the BF, and implanted optical fiber in BF. Left bottom, average traces with standard deviation show postsynaptic activity with 30min photostimulation at 0, 3, 5, and 10Hz 10ms pw. Right top, plot shows maximum postsynaptic activity in relation to stimulation frequency (N=6). Right bottom, plot shows postsynaptic activity at baseline (1min before stimulation), early stimulation (1-5min), and late stimulation (25-30min) at different frequencies (N=6). n.s. not significant. **d.** Left top, schematic shows Chrimson in LC^NE^ neurons, GCaMP6f in the Thal, and implanted optical fiber in Thal. Left bottom, average traces with standard deviation show postsynaptic activity with 30min photostimulation at 0, 5, and 10Hz 10ms pw. Right top, plot shows maximum postsynaptic activity in relation to stimulation frequency (N=6). Right bottom, plot shows postsynaptic activity at the baseline (1min before stimulation), early stimulation (1-5min), and late stimulation (25-30min) at different frequencies (N=6). n.s. not significant. **e.** Left, schematic shows GCaMP6s in LC^NE^ neurons, GRAB_NE or GCaMP6f in the BF, and tail suspension. Right, plots show corresponding traces of the LC^NE^ activity (N=6), BF NE release (N=8), and BF postsynaptic activity (N=6), and corresponding AUC in 1 min binning normalized to 1 min before 10min tail suspension. **f.** Left, schematic shows forced swim. Right, plots show traces of the LC^NE^ activity (N=7), BF NE release (N=6), and BF postsynaptic activity (N=7); and corresponding AUC in 1 min binning normalized to 1 min before forced swim.

Given the observed NE depletion and postsynaptic desensitization in BF with photostimulation, we then asked whether these effects occur under naturalistic environmental stimulation such as during acute stress exposure. Cohorts expressing either GRAB_NE or GCaMP6f in the BF were used to determine axonal release and postsynaptic activity, respectively, while another cohort expressing GCaMP6s in LC^NE^ neurons of Dbh-cre mice was used to determine LC^NE^ activity. We first tested tail suspension; and observed that the LC^NE^ activity remained continuously elevated during a long tail suspension session, with the NE release steadily increased and appeared to plateau, and the postsynaptic activity initially increased then gradually decreased (**Fig 2e**). Pre-treating with 10mg/kg prazosin, but not 10mg/kg propranolol or saline, caused a faster decay in postsynaptic activity (**SFig 2b**). Together, these results indicate that although tail suspension does not deplete axonal NE, the postsynaptic activity decreases over time, likely due to well-established receptor desensitization of adrenergic receptors^26^. We then tested forced swim, an established acute stressor in rodents^27^, and observed LC^NE^ activity initially increased, then decreased sharply after 1 min. Interestingly, NE release again monotonically increased, though at a slower rate than tail suspension, while the postsynaptic activity reflected LC^NE^ activity with an initial increase, followed by a decrease after 1 min (**Fig 2f).** These results show that NE release does not acutely deplete, but a decreased postsynaptic response is observed on the order of minutes.

We then sought to determine whether NE depletion from increased tonic LC^NE^ activity affects the phasic mode of LC^NE^ transmission. Using ChrimsonR expressed in the LC^NE^ and GRAB_NE in the BF, we generated a range of tonic LC^NE^ activity by tonically photostimulating the LC^BF^ terminals at 0, 3, 5, and 10Hz for 30min. Concurrently, we simulated regular phasic activity (typically ∼10-20Hz, 200-500ms for LC^12^) by photostimulating at 20Hz, 10ms pw for 200ms every 20s. As seen in **Fig 3a**, the progressive increased NE release from higher tonic stimulation frequencies is associated with a significant decreased release from phasic stimulations. This inverse relationship suggests that both transmission modes draw from the same NE pools. Additionally, the decreased phasic response is not due to α2-mediated inhibition of release, since we found that pre-treating with atipamezole did not prevent this decrease (**Fig 3b**). In a complementary experiment, we further examined simulated phasic NE release during changes in tonic activity from acute stressors. During a 10min tail suspension, for instance, tonic NE release remained continually elevated, resulting in a significant decrease in phasic NE release as compared to baseline (**Fig 1c**). However, during restraint stress, NE release was only transiently elevated, and the simulated phasic NE release during restraint did not significantly differ from before (**SFig 2c**). We also stressed the animals with recurrent footshocks to elicit high, but periodic increases in phasic NE release as shown in **Fig 3d**. Interestingly, after the initial footshock, the basal NE release increased and remained elevated for the duration of the session. The phasic NE release from repeated footshocks, on the other hand, diminished over the course of the session, indicating that increased NE release from tonic activity limits the release from phasic activity. This decreased phasic NE release was not due to decreasing LC^NE^ activity over the session (**Fig 3d**), nor due α2-mediated inhibition of LC^NE^ activity or NE release (**SFig 2d,e**). Together, these experiments demonstrate that phasic NE release can be directly influenced by the extent of tonic NE release depending on stressor type or exposure.

**Figure 3.**
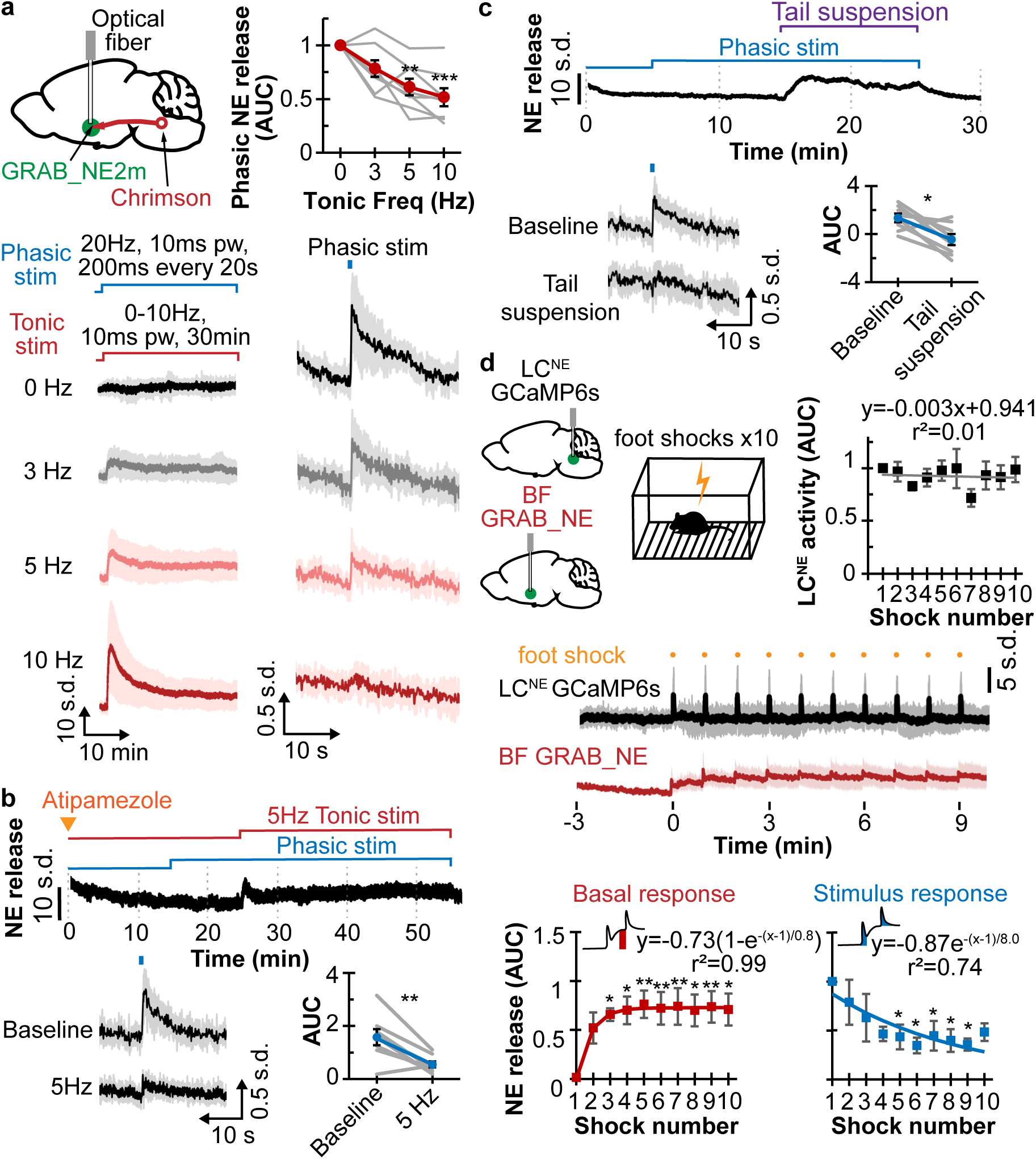
**a.** Top left, schematic shows Chrimson expressed in the LC^NE^ neurons, GRAB_NE2m in the BF, and an implanted optical fiber in BF. Bottom, traces of NE release for 30min of 0, 3, 5, and 10Hz tonic photostimulation (left) with 20Hz, 10ms pw, 200ms phasic stimulation every 20s (right). Top right, plot shows AUC of phasic NE release in relation to differing tonic stimulation frequencies (N=8). **p<0.01, ***p<0.001. **b.** Top, trace shows NE release in response to 5Hz tonic photostimulation with 20Hz, 10ms pw, 200ms phasic stimulation 20min after 1mg/kg atipamezole treatment. Bottom left, averaged traces in response to phasic stimulation at baseline and 5Hz tonic stimulation. Bottom right, corresponding AUC of phasic NE release is shown in plot (N=8). **p<0.01. **c.** Top, trace shows NE release in response to 10min tail suspension with 20Hz, 10ms pw, 200ms phasic stimulation. Bottom left, averaged traces in response to phasic stimulation at baseline and during tail suspension. Bottom right, corresponding AUC of phasic NE release is shown in the plot (N=8). *p<0.05. **d.** Top left, schematic shows GCaMP6s in the LC^NE^ neurons, GRAB_NE in the BF, and mouse experiencing 10 footshocks. Middle, average traces with standard deviation of LC^NE^ activity (black) and GRAB_NE in the BF (red) in response to footshocks. Top right, plot shows the AUC of LC^NE^ activity in response to the footshocks (N=7). Bottom, plots show the basal changes to NE release (left, red) and AUC of NE release (right, blue) in response to footshocks (N=8). Exponential fitting curves and equations are also shown. *p<0.05, **p<0.01.

Next, we determined whether dopamine (DA), another catecholamine, has similar constraints on release and on modes of transmission (tonic ∼4-5Hz, phasic ∼20Hz, <200ms)^13, 14^; and whether these features we identified for the LC^NE^ release are similar in the dopamine system. ChrimsonR was expressed in DA neurons in the ventral tegmental area (VTA) of DAT-cre mice; a fluorescent DA sensor GRAB_DA2m was expressed in the nucleus accumbens core (NAc), with a fiber optic placed in the NAc (**SFig 3a**). When repeated axonal photostimulations were applied for 1min at 10Hz, 50ms pw, DA release was robustly elicited, with decreased S_1_ (72±7%) and S_2_ (69±6%) responses. Increasing stimulation frequency to 20Hz, 10ms pw, showed similar decrease in S_1_ (77±6%) and S_2_ (75±4%). Photostimulating for 30min at 0, 5, and 10Hz 5ms pw (**SFig 3b**) also produced increased DA release with increased stimulation frequency, without significant decrease in the late part of the stimulation. Using the same 10ms pw as the NE stimulation exhibited similar trend as the 5ms pw at 10Hz (**SFig 3c**). Furthermore, when simulated 200ms phasic stimulations at 20Hz 5ms pw every 20s were added to these various tonic stimulation frequencies, the phasic DA release decreased as the tonic frequencies increased (**SFig 3d**). These experiments show that although NAc DA also show signs of depletion at the high tonic activity, the DA pools replete more rapidly than the NE pools we examined in the BF and Thal. Nevertheless, taken together, these results demonstrate an optimal activity window for catecholamine signaling at the synaptic level.

## Discussion

In this study, we used two modern biosensor approaches to unveil for the first time two distinct pools of the respective catecholamines NE and DA based on fast and slow repletion kinetics *in vivo* (**Fig 1C**). The fast-repleting pool constitutes ∼25-50% of NE transmission in the brain regions examined from our axonal photoactivation, but constitutes a much larger proportion, ∼70-75%, of DA transmission. For NE, the slow repletion takes place on the order of tens of minutes, suggesting that these different pools may translate to differing time scales on which these neuromodulators can act to regulate behavior.

Interestingly, the NE fast repleting pool strongly depends on NET (**Fig 1D**), which previously has been shown to colocalize with Rab11, a marker of recycling endosomes^28^, suggesting a potential source for this NE pool. However, it is important to consider the several other possibilities for these different pools. NE and DA are reported to reside in both small synaptic vesicles and large dense core vesicles^29–31^, and the differing molecular compositions of these vesicles that are thought to explain exocytic differences^32^, may explain endocytic differences as well in repletion. These pools may also differ in their synaptic localization as readily releasable versus reserve pools^33–35^. They may also represent differing release from different axons, as seen before with NE release^36^ or represent (synaptic) point-to-point versus (non-synaptic) volumetric transmission^37, 38^. Thus, the molecular identities of these pools remain to be seen and an important area for further exploration using modern omics approaches. The measured NE depletion from the same stimulus also differed depending on the brain region, likely due to variations in NET and α2-AR levels as observed across LC^NE^ subpopulations from single-nuclei RNA sequencing^24^, and suggests that region-specific NE availability may help to explain the non-homogenous effect that increased LC^NE^ activity has across the brain^39^. Additionally, given recent suggestion that NE and DA may be co-released from the LC^40–42^, together with our observation of a steep NE depletion in the BF and Thal (**Fig 1E**) as compared to the DA depletion in the NAc (**SFig 3B**), it would be important to determine whether co-released NE depletes more rapidly than DA within the same brain regions.

Furthermore, our observation that NE and DA gradually exhaust their release pools at higher stimulation frequencies, mirrored by a gradual decrease in postsynaptic calcium, provides an overdue plausible mechanistic explanation for the inverted-U relation. In particular, as phasic activity of LC^NE^ is associated with focus task performance^8^, our finding that higher tonic NE release from increased tonic activity causes a proportional reduction in NE output from phasic activity provides a natural explanation for the decreased behavioral performance at the higher LC^NE^ activity. Interestingly, tail suspension and force swim, two highly stressful, acute interventions known to increase LC^NE^ firing^34^ did not deplete NE, but did exhibit a gradual decrease in postsynaptic calcium following an acute rise, suggesting that acute stressors do not exhaust brain NE, likely due to negative feedback or asynchronous release kinetics. The postsynaptic calcium response, on the other hand, diminishes in minutes, likely due to desensitization of the α_1_-ARs^43, 44^, though downstream signaling may still persist^45^. The temporal pattern of β-AR-mediated signaling, however, remains to be explored in these scenarios, and will require molecular tools, like CRISPR to isolate discrete subtype selective combinations in given molecularly identified post-synaptic neurons. Thus, postsynaptic AR-desensitization appears to be the predominant mechanism in acute stress, and catecholamine depletion may play a more important role in more prolonged elevation in neuronal firing.

Moreover, we show that the tonic and phasic LC and DA firing, as two modes of transmission, exhibit an inverse relationship (**Fig 3A, SFig 3D**), suggesting they draw from the same respective catecholamine pools. In particular, in situations where LC tonic activity is high, such as during certain acutely stressful events, the NE signal-to-noise from phasic firing is greatly diminished, and thus, contribute to increased exploratory behaviors and decreased task performance^8, 46^. NE levels from prolonged elevated LC activity from chronic stress^47^ remains to be explored, as possible NE depletion and the brain’s normal versus impaired compensation may have implications in psychiatric disorders and treatments. Additionally, the study further corroborates the now well-hypothesized impairment of phasic activity from increased tonic activity that may play an explanatory role in neurodegeneration in Alzheimer’s and Parkinson’s diseases^48^, especially given the modulatory role LC has on brain waste clearance^49, 50^.

In summary, this study demonstrates the *in vivo* constraints on catecholamine signaling at the higher ends of neural activity, showing catecholamine depletion at the axons, desensitization at the postsynapse, and overall decreased signal-to-noise in the phasic mode of transmission. These constraints introduce a form of non-linearity in neuromodulation occurring at level of synaptic transmission that may help explain the upper end of the inverted-U relationships between catecholaminergic neural activity and cognitive performance^7, 8, 11^, and contribute to greater insights into catecholamine-associated psychiatric disorders such as depression and substance use disorder,^47, 51–54^ and neurodegenerative diseases^55, 56^.

## Methods

### Animals and surgery

Adult male and female Dbh-cre C57BL/6J mice (Dbh*^tm3.2(cre)Pjen^*, Jackson Laboratory #033951) were group housed on a 12:12 h reverse light-dark cycle and given food and water *ad libitum*. Surgeries were performed on mice ages >8 weeks under 1.5-2.0% isoflurane. Mice were injected in the LC (AP −5.45mm, ML +1.1mm, DV - 4.2 to −3.7mm DV to bregma) using a medial-facing beveled 1 µl Hamilton syringe. A blunt tip 1 µl Hamilton syringe was used for injections to the posterior basal forebrain (BF) (AP +0.1mm, ML +1.3mm, DV −5.3 to −5.15mm), the medial thalamus (mThal) (AP −1.7mm, ML +1.0mm at 15° angle, DV −3.95 to −3.8mm), the ventral tegmental area (VTA) (AP - 3.4mm, ML +0.6mm, DV −4.7 to −4.4mm), and nucleus accumbens core (NAc) (AP +1.4mm, ML +1.5mm at 10° angle, DV −4.1 to −3.8mm). Respective 400 µm optical fibers (Doric Inc., MFC_400/430-0.48_MF2.5_FLT) were placed along the same AP and ML coordinates except DV is −3.8mm for LC, −5.2mm for BF, −3.8mm for mThal, −3.85mm for NAc and secured using Metabond (Parkell #S380). Mice recovered for at least 5 weeks before experiments to allow optimal viral expression. All experiments are in accordance to IACUC protocol.

Viruses
|  |  |
| --- | --- |
| AAV5-EF1α-DIO-EYFP | Addgene #20756 |
| AAV5-Syn-FLEX-rc[ChrimsonR-tdTomato] | Addgene #62723 |
| AAV-DJ-EF1α-DIO-GCaMP6s | Stanford Gene Vector and Viral Core |
| AAV-DJ-EF1α-DIO-GCaMP6f | Stanford Gene Vector and Viral Core |
| AAV-hSyn-GRAB_NE2m | Yulong Li (Peking University) |
| AAV-hSyn-GRAB_NE2h | Yulong Li (Peking University) |
| AAV9-hSyn-GRAB_DA2m | Addgene #140553 |

### Immunohistochemistry

Immunohistochemistry (IHC) was performed as described previously (McCall *et al*., 2017). Briefly, mice were euthanized with sodium pentobarbital and transcardially perfused with 4% paraformaldehyde (PFA), post-fixed for 1-3 days in 4% PFA, and then cryo-protected in 30% sucrose. Brain sections (30-100µm) were collected and kept in 0.1M PB at 4°C until IHC. Sections were washed 3 times for 10min in PBS, then 1h in blocking buffer (0.5% Triton X-100 and 5% goat serum in 0.1M PBS) at room temperature, followed by overnight incubation with primary antibody () at 4°C. Next, the sections were again washed 3 times for 10min in PBS, incubated for 1h in Alexa Fluor secondary antibodies (1:1000) at room temperature, and washed 3 times for 10min in PBS. Then, the sections were mounted with VectaShield Vibrance Hardset mounting medium (Vector Laboratories) with DAPI, and coverslips placed. Images were acquired on a epifluorescent microscope (Leica DFC700T).

### Fiber photometry and photostimulation

Fiber photometry recordings of the genetically encoded fluorescent sensors were performed as previously described. Briefly, the implanted fiber was connected to the patch cable via a ferrule sleeve. A real-time processor (TDT RZ5P, sampling rate of 1017.25Hz) recorded the filtered emission (Doric FMC4 filter cube) from the fluorescent sensor at upon excitation at 470nm and 405nm using the accompanied TDT Synapse software. For terminal stimulations, the same photometry fiber served as conduit for 565nm LED photostimulation adjusted for light intensity ∼2.5µW at the optical fiber tip.

Photometry recordings were analyzed using custom python script. Baseline drift from photobleaching artifact was corrected with an exponential decay curve. For GRAB_NE and GCaMP signals, dF/F was calculated as the linear least squared fit of 405nm signals subtracted from the 470nm signals. The 405nm signals were not used in GRAB_DA as it does not appear to serve as an appropriate isobestic channel. Z-scores were calculated using the mean and standard deviation of a 5-10 min baseline before photostimulation.

### Drugs

All drugs were administered intraperitoneally (*i*.*p*.). These include reserpine (Sigma 83580) dissolved in 0.2% glacial acetic acid to 0.5 mg/ml; nepicastat (Sigma-Aldrich SML0940) suspended as 5mg/ml in saline mixed with 1.5% DMSO, 1.5% Kolliphor; dexmedetomidine (100 mcg/ml Piramal Critical Care, PSLAB-020872-00) diluted to 5mcg/ml in saline, atipazemole made to 0.1mg/ml in saline, atomoxetine hycrochloride (thermoscientific, Cat 467680010) make to 0.3mg/ml in saline, propranolol hydrochloride (Tocris Bioscience Cat. 0624) made to 1mg/ml in sterile water, prazosin hydrochloride (Tocris Bioscience Cat. 0623) made to 1mg/ml in distilled water. Drugs were administered at least 15-20 min prior to experiments unless otherwise noted.

### Stress behavioral assays

#### Tail suspension

Mice were suspended by an experimenter holding the tail for 10 min ∼10in above the floor.

#### Forced swim

Mice were transferred from a holding chamber into a 5L cylindrical container containing 4L of water (∼24°C) and allowed to swim up to 15min. Mice were removed from water if there was evidence of drowning. Mice were subsequently dried on a heating pad for recovery.

#### Physical restraint

Mice were placed in a conical tube with a strip cut open on top to allow photometry patch cable attachment and the front tapered end cut open to allow nose access. The other end has a lid with a center hole for the tail.

#### Foot shock

Mice were placed in a Med Associates Fear Conditioning Chamber (NIR-022MD, 29.53cm L x 23.5 cm W x 20.96cm H) with a conductive grid floor, a soundproof barrier, and an infrared lighting. They were habituated to the box for 3 min before experiencing 1s 0.5mA shocks every min for 10 minutes.

### Statistical analysis

All summary data are expressed as mean±SEM. Statistical significance was denoted as *p<0.05, **p<0.01, ***p<0.001, ****p<0.001 as determined by Student’s t-test, one-way or two-way repeated measure analysis of variance (ANOVA), followed by Dunnett’s post-hoc test. Statistical tests were performed in Excel and in R, and a summary of the statistical tests performed and p-values are shown in **Supplemental Table 1**.

## Data and code availability

Custom python analysis code, and photometry data will be provided upon request.

## Acknowledgements

We thank all of Bruchas lab for their helpful insights and discussions with manuscript. We acknowledge our funding sources; FAER MRTG (L.L.), K99/R00 DA053336 (L.L.), Mary Gates Research Scholarship (E.M.L.), National Natural Science Foundation of China grants 31925017 and 31871087 (Y.L.), and R01 MH112355 (M.R.B).

## Authors contributions

L.L. A.R. and E.M.L. collected data. J.F. and Y.L. provided reagents. L.L. and M.R.B. designed experiments, analyzed data, and wrote the paper.

---

**Supplemental Figure 1.**
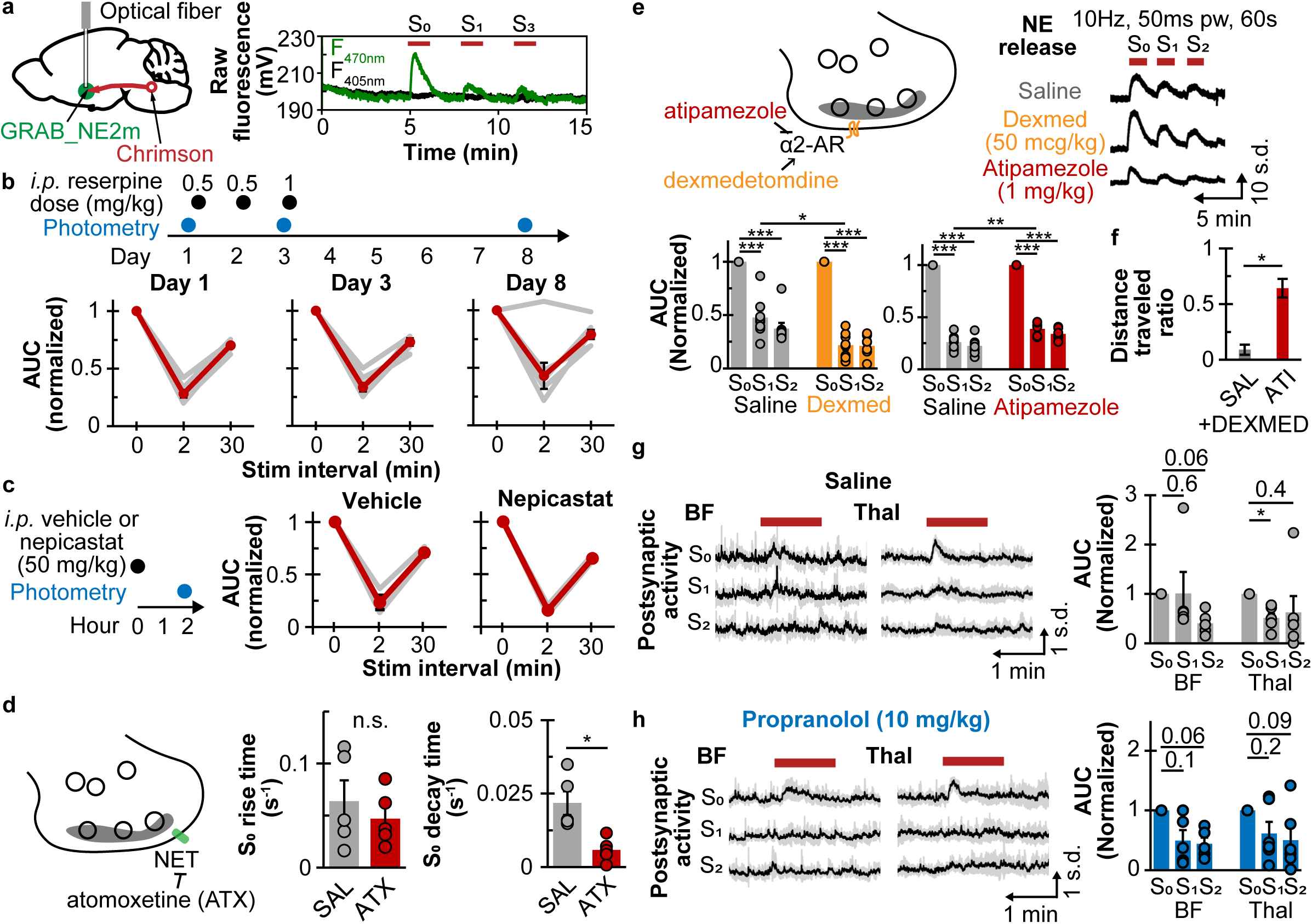
**a.** Schematic (left) shows Chrimson in the LC^NE^ neurons, GRAB_NE2m in the BF, and implanted optical fiber in BF. Raw traces (right) show GRAB_NE2m fluorescence at 405nm (black) and 470nm (green) in response to 10Hz photostimulation. **b.** Top, time line shows repeated *i.p.* reserpine treatments and photometry recordings. Bottom, plots of AUC of repeated 10Hz (50ms pw) photostimulation normalized to the first stimulation at 2 or 30min intervals on day 1, 3, and 8 during reserpine treatments. **c.** Left, timeline shows either *i.p.* vehicle or 50mg/kg nepicastat treatments and photometry recordings. Right, plots of AUC of repeated 10Hz (50ms pw) photostimulation normalized to the first stimulation at 2 or 30min intervals for vehicle and nepicastat. **d.** Left, schematic illustrates atomoxetine (ATX) inhibition of norepinephrine reuptake transporter (NET). Right, plots show the rise (left) and decay (right) constant of NE release to the first stimulation with either *i.p.* saline (SAL) or 3mg/kg atomoxetine treatments (N=5). *p<0.05. n.s. not significant. **e.** Top left, schematic illustrates α2 adrenergic receptor activation by dexmedetomidine (DEXMED) and inhibition by atipamezole. Top right, representative traces show NE release in response to repeated 10Hz photostimulation 20min after *i.p.* saline, 50mcg/kg dexmed, and 1mg/kg atipamezole. Bottom, plots show the corresponding AUC normalized to the first stimulation for these treatments (N=8). *p<0.05, **p<0.01, ***p<0.001. **f.** Plot shows the ratio of distance traveled for mice pre-treated with either *i.p.* saline (SAL) or 1mg/kg atipamezole (ATI) before and after 50mcg/kg dexmed administration (N=4). *p<0.05. **g.** Left, average traces with standard deviation show the postsynaptic activity in the BF and Thal in response to the repeated 10Hz photostimulations 20min after *i.p.* saline. Right, plots of the corresponding AUC normalized to the first stimulation in BF (N=5) and Thal (N=6). *p<0.05. **h.** Similar plots as in g except mice are treated with 10mg/kg propranolol.

**Supplemental Figure 2.**
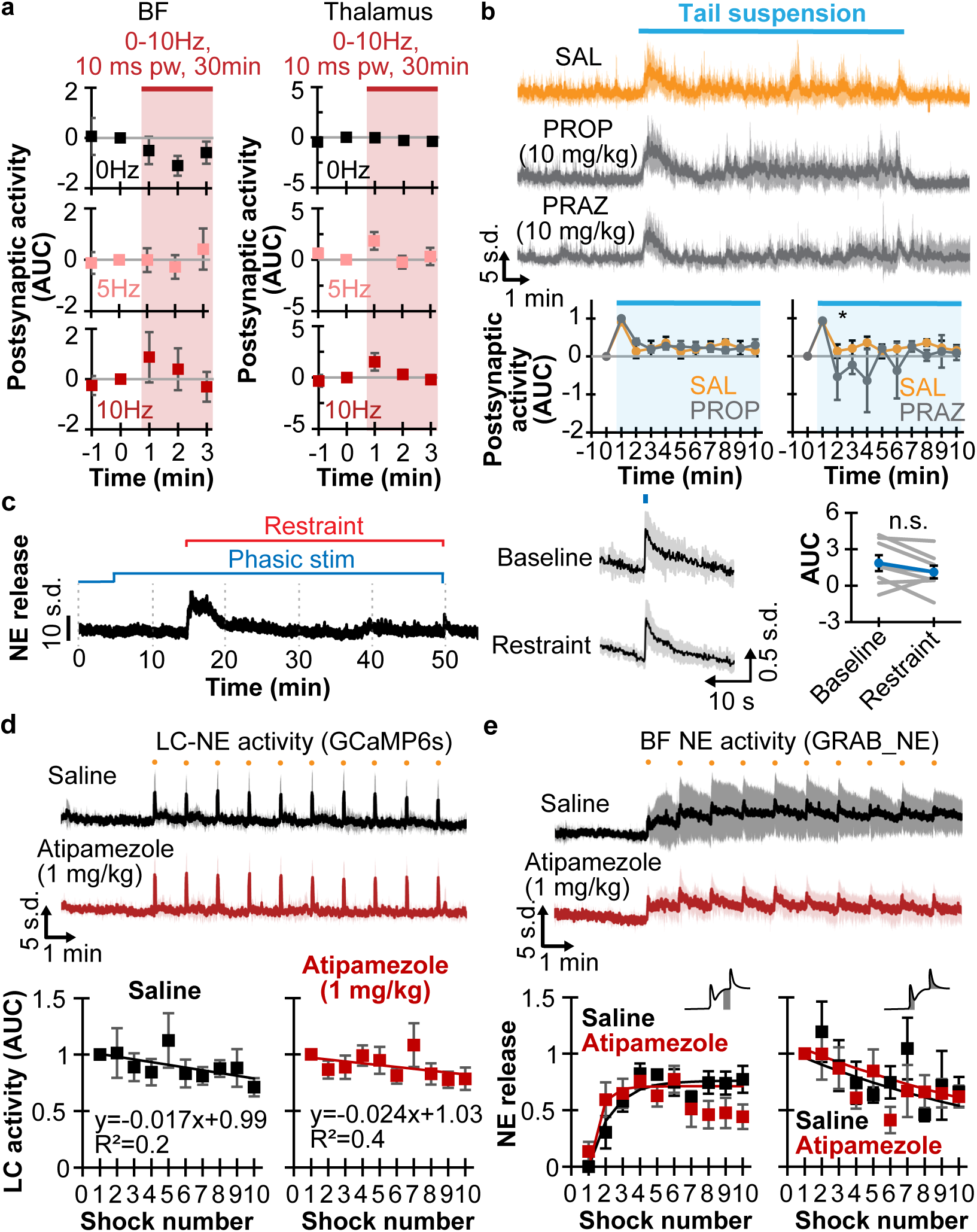
**a.** Plots show the AUC of postsynaptic activity in 1min binning and normalized to 1 min before 0, 5, and 10Hz photostimulation in the BF (left, N=6) and thalamus (right, N=6). Red bar and shading show stimulation. **b.** Top, traces show BF postsynaptic activity in response to 10min tail suspension 20min after *i.p.* saline (SAL, orange), 10m/kg propranolol (PROP), and 10mg/kg prazosin (PRAZ). Bottom, plots shown AUC of normalized postsynaptic activity to baseline and peak activity in 1 min binning comparing SAL and PROP; and SAL and PRAZ, respectively. *p<0.05 using K-S test. **c.** Left, photometry trace shows NE release in response to physical restraint and phasic photostimulations 20Hz, 10ms pw, 200ms every 20s. Middle, averaged photometry trace in response to phasic stimulation at baseline and during restraint. Right, plot shows AUC of phasic response at baseline and restraint (N=8). n.s. not significant. **d.** Top, photometry traces show LC^NE^ activity after *i.p.* saline (black) and 1mg/kg atipamezole (red) in response to 10x footshocks. Bottom, plots show corresponding AUC of LC^NE^ activity to footshocks with saline and atipamezole treatments (N=7). **e.** Top, photometry traces show BF NE activity after *i.p.* saline (black) and 1mg/kg atipamezole (red) in response to 10x footshocks. Bottom, plots show corresponding AUC of NE release to footshocks with saline and atipamezole treatments (N=6).

**Supplemental Figure 3.**
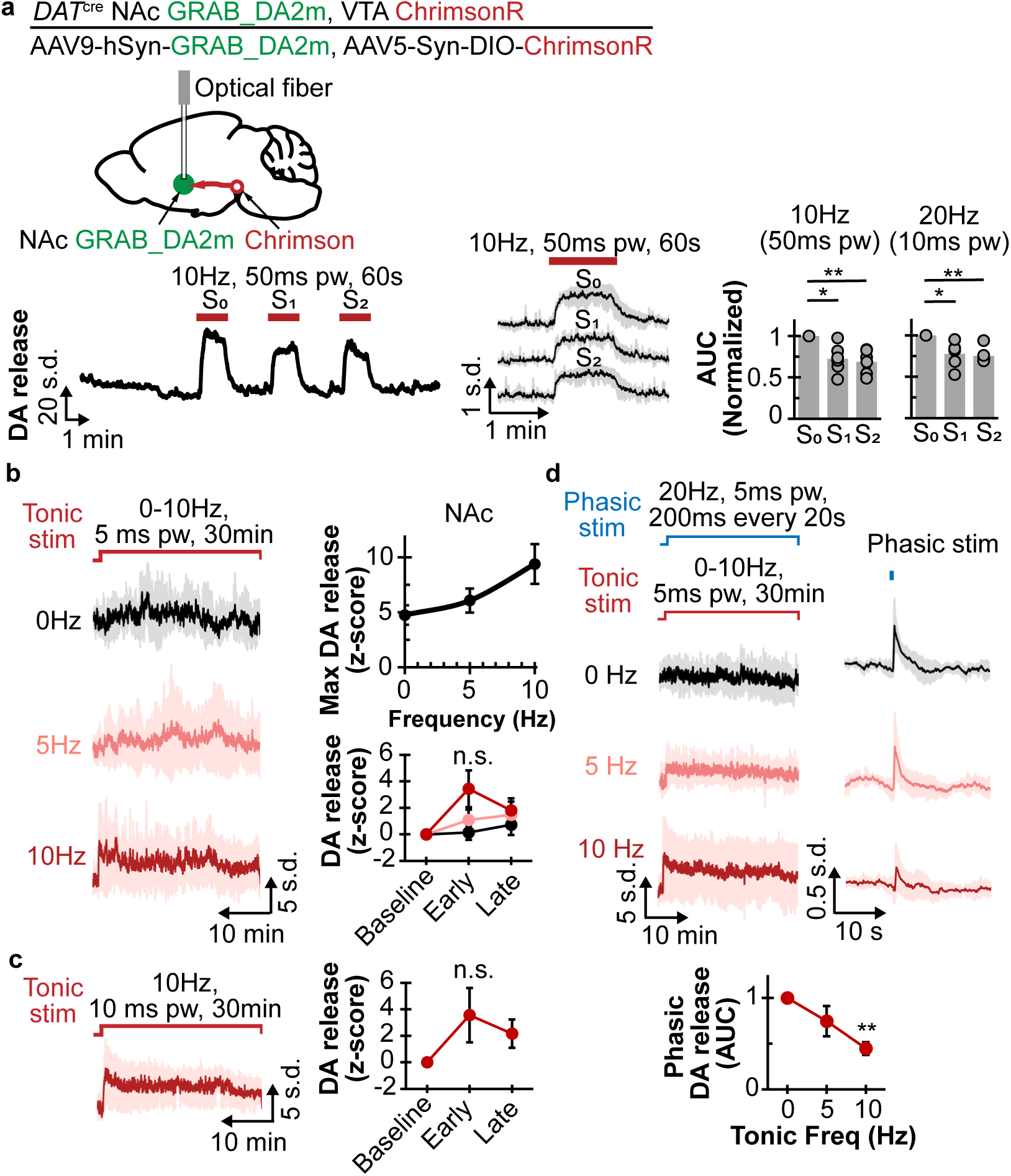
**a.** Top, schematic shows Chrimson expressed in the ventral tegmental area (VTA) in DAT-cre mice, GRAB_DA2m in the nucleus accumbens core (NAc), and implanted optical fiber in the NAc. Bottom left and middle, photometry trace of dopamine (DA) release in response to repeated 10Hz (50ms) photostimulation. Bottom right, plots show AUC normalized to the first stimulation using 10Hz, 50ms pw, and 20Hz, 10ms pw (N=6). *p<0.05, **p<0.01. **b.** Left, photometry traces of DA release with 30min of photostimulation at 0, 5, and 10Hz 5ms pw. Right top, plot shows maximum DA release in relation to tonic frequency (N=6). Right bottom, plot shows DA release normalized to baseline (1 min before stimulation), early (1-5min after stimulation), and late (25-30min after stimulation) for the different stimulation frequencies (N=6). n.s. is not significant. **c.** Left, photometry trace of DA release in response to 30min photostimulation at 10Hz, 10ms pw. Right, plot shows AUC of DA release at baseline (1 min before stimulation), early (1-5min), and late (25-30min) for 3 mice. n.s. not significant. **d.** Top, photometry traces of DA release with 30min of tonic photostimulation at 0, 5, and 10Hz 5ms pw (left) and 200ms phasic stimulation at 20Hz 5ms pw every 20s (right). Bottom, plot shows AUC of phasic photostimulation in relation to tonic stimulation frequency (N=6). **p<0.05.

